# Calcium activates purified human TRPA1 with and without its N-terminal ankyrin repeat domain in the absence of calmodulin

**DOI:** 10.1101/2020.03.19.999276

**Authors:** Lavanya Moparthi, Satish Babu Moparthi, Jérôme Wenger, Peter M. Zygmunt

## Abstract

Extracellular influx of calcium or release of calcium from intracellular stores have been shown to activate mammalian TRPA1 as well as to sensitize and desensitize TRPA1 electrophilic activation. Calcium binding sites on both intracellular N- and C-termini have been proposed. Here, we demonstrate based on fluorescence correlation spectroscopy (FCS), Förster resonance energy transfer (FRET) and bilayer patch-clamp studies, a direct calmodulin-independent action of calcium on the purified human TRPA1 (hTRPA1), causing structural changes and activation of hTRPA1 with and without its N-terminal ankyrin repeat domain (N-ARD). Thus, calcium can activate hTRPA1 by direct interaction with binding sites outside the N-ARD.

## Introduction

The mammalian pain receptor TRPA1 was initially identified as a noxious cold sensor and chemoreceptor of electrophilic irritants as well as non-electrophilic cannabinoids (1-3). Importantly, TRPA1 was also shown to be activated by an increase in the intracellular free calcium concentration (2). Whereas electrophiles are believed to promote TRPA1 channel opening primarily by binding to cysteine residues within the cytoplasmic N-terminus, the mechanism by which calcium modulates TRPA1 activity is not obvious (4, 5). Studies have shown a direct activation of TRPA1 by calcium in cell-membrane inside-out patches containing heterologously expressed TRPA1 (6-8). However, an effect solely by calcium has been questioned as membrane bound calcium co-mediators such as calmodulin may still be present in isolated membrane patches (9). Some studies have focused on the modulatory role of calcium on electrophilic TRPA1 activation, and also with regard to calmodulin dependence (9-13). Furthermore, structures within the intracellular N- and C-termini have been suggested to contain calcium binding sites coupled to the gating of TRPA1 (7-9, 11, 13). Interestingly, a calcium binding amino acid cluster has been identified in the C-terminus, and recently in the cytoplasmic end of transmembrane domain 2 and 3 of TRPA1 (11, 13, 14).

In this study, we asked if calcium alone modifies the structure and induces activation of the purified hTRPA1 and Δ1-688 hTRPA1 without the N-terminal ankyrin repeat domain (N-ARD) (Fig. 1 A). We report, based on fluorescence correlation spectroscopy (FCS), Förster resonance energy transfer (FRET) and bilayer patch-clamp studies of purified hTRPA1 with and without its N-ARD, a direct non-calmodulin-dependent action of calcium on hTRPA1, causing structural changes and hTRPA1 channel activity.

**Figure 1.**
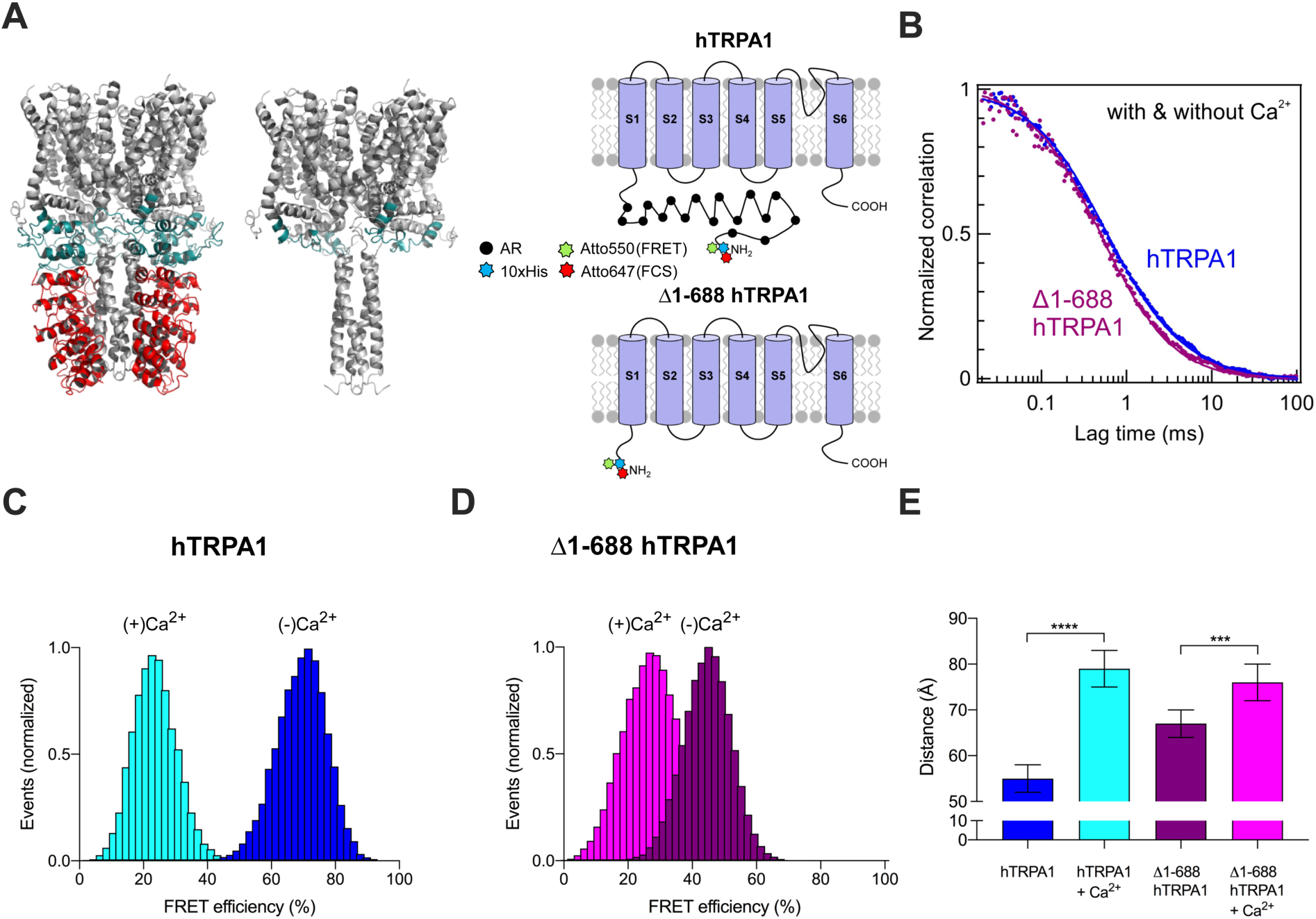
Structural rearrangements of hTRPA1 with and without its N-terminal ARD (Δ1-688 hTRPA1) by calcium. (A) Tetrameric structure of hTRPA1 with and without its N-terminal ankyrin repeat domain (N-ARD) highlighted in red; Protein Data Bank (PDB) accession number 3J9P. Schematic representation of the full length hTRPA1 monomer and truncated Δ1-688 hTRPA1 monomer lacking ankyrin repeats (AR). The N-terminus His-tag was labeled with the fluorophore Atto647N and Atto550 in FCS and FRET experiments, respectively. (B) Normalised Fluorescence Correlation Spectroscopy (FCS) traces of hTRPA1 and Δ1-688 hTRPA1. Similar traces were obtained in presence and absence of 2 mM calcium, indicating that calcium did not noticeably modify the hydrodynamic radius monitored in FCS. (C-E) Single-molecule Förster Resonance Energy Transfer (FRET) efficiency histograms of intramolecular fluorophore dual-labeled (N-His-tag = Atto550 and lysines = Atto647N) hTRPA1 (C) and Δ1-688 hTRPA1 (D) in the absence and presence 200 µM CaCl_2_. Gaussian and lognormal distributions are used to fit the data, and a higher value in FRET histograms indicates a higher degree of compactness of the protein. (E) Calculated average donor-acceptor intramolecular distance in ångström (Å) obtained from the FRET data in C and D. Data is the mean ± SD of 5 (B) and 3 (E) separate experiments. \*\*\**P* < 0.001 and *\*\*\*\*P* < 0.0001, Student’s unpaired t-test. All experiments were performed at 22°C after pre-incubation with CaCl_2_ for 2 h at 22°C.

## Results

### Effect of calcium on hTRPA1 structure analyzed by FCS and FRET

Fluorescence Correlation Spectroscopy (FCS) monitors the average diffusion time of the protein across a fixed femtoliter detection volume. From the FCS data, the diffusion coefficient and the hydrodynamic radius can be obtained from the Stokes-Einstein equation. The FCS data recorded for hTRPA1 and Δ1-688 hTRPA1 indicate monophasic correlation curves with diffusion times of 560 ± 40 µs (D = 3.6 × 10^−7^ cm^2^·s^−1^) and 500 ± 30 µs (D = 4.1 × 10^−7^ cm^2^·s^−1^), respectively, in aqueous solution (Fig. 1B). The corresponding hydrodynamic radii are 6.2 nm for hTRPA1 and 5.6 nm for Δ1-688 hTRPA1. Faster diffusion time and smaller hydrodynamic radius for the truncated Δ1-688 hTRPA1 as compared to the full length hTRPA1 is in line with the lack of the N-terminal ankyrin repeat domain for the truncated protein. However, the FCS data did not reveal any major change in the presence or absence of calcium for both protein systems. We relate this effect to the fact that FCS is only sensitive to the average hydrodynamic radius of the system and lacks more accurate resolution on the protein conformation.

To gain more insight in the possible structural changes induced by calcium, we use intramolecular FRET between donor and acceptor dyes labelled within the protein. The effect of calcium on hTRPA1 and Δ1-688 hTRPA1 is clearly evidenced in the FRET energy transfer efficiency (*E*) histograms (Fig. 1 C,D). In the absence of calcium, hTRPA1 displayed an average FRET efficiency *E* = 0.72, equivalent to a global distance of 55 ± 5 Å between the intramolecular donor and acceptor fluorophores (Fig. 1 C,E). In the presence of calcium, hTRPA1 changed its structure significantly as indicated by *E* = 0.24, corresponding to a global distance of 79 ± 5 Å between the donor and acceptor fluorophores (Fig. 1 C,E). Likewise, calcium increased the global distance between the Δ1-688 hTRPA1 intramolecular donor and acceptor fluorophores; *E* = 0.45 and at *E* = 0.28, in the absence and presence of calcium, respectively, corresponding to a calcium-evoked increase in global distance from 67 ± 5 Å to 76 ± 5 Å (Fig. 1 D,E). Importantly, the fraction of samples containing only the donor dye was negligible in these experiments, as illustrated by the absence of only-donor peak centered at *E* = 0.

### Effect of calcium on hTRPA1 channel activity

The purified hTRPA1 and Δ1-688 hTRPA1 were reconstituted into artificial lipid bilayers for electrophysiological recordings of channel activity. Both hTRPA1 and Δ1-688 hTRPA1 responded to calcium with vivid channel activity (Fig. 2 A-C) including variability in the single-channel open probability (Fig. 2D). Several single-channel current levels were observed at both 10 and 100 μM (Fig. 2 A-C), also reflected by the variation in single-channel conductance values (Fig. 2D). The activity of hTRPA1 at 10 and 100 μM was abolished by the TRPA1 antagonists A-967079 and HC030031 (Fig. 2C), both obtained from Sigma-Aldrich. A-967079 inhibits hTRPA1 with an IC_50_ value of 67 nM and is about 100 times more potent than HC030031 (15, 16), which was used in this study at a concentration that abolished purified hTRPA1 and Δ1-688 hTRPA1 single-channel activity evoked by chemical ligands, temperature and mechanical stimuli (17-20). As also reported previously (17-20), no basal TRPA1 activity was observed at 20-22°C (not shown). Likewise, no channel activity was observed in membranes without the purified hTRPA1 proteins exposed to 10 and 100 μM calcium (n=3, not shown).

**Figure 2.**
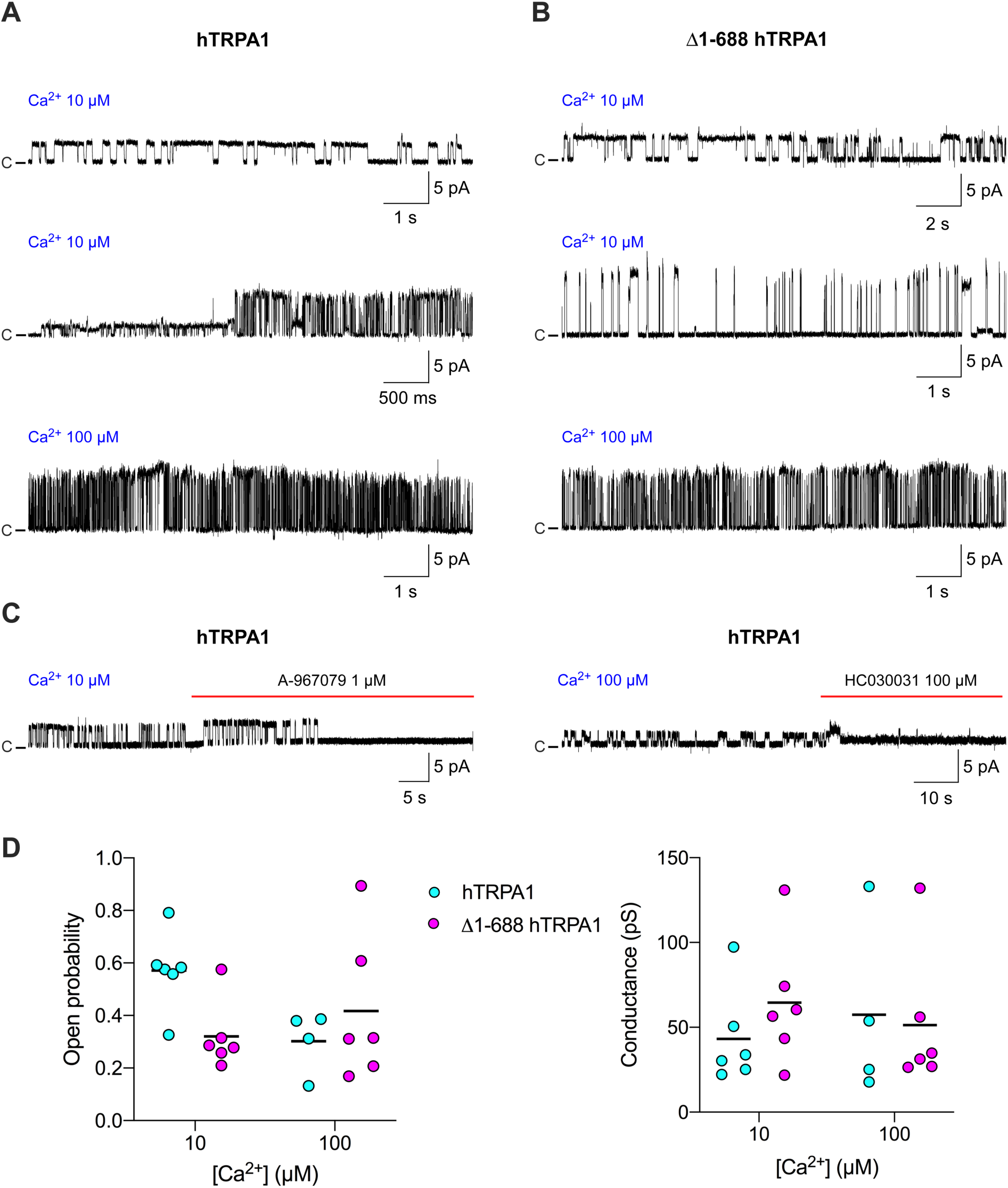
Activation of human TRPA1 with and without its N-terminal ARD (Δ1-688 hTRPA1) by calcium. Purified hTRPA1 and Δ1-688 hTRPA1 were reconstituted into planar lipid bilayers and single-channel currents were recorded with the patch-clamp technique in a symmetrical K^+^ solution at a test potential of +60 mV (20-22°C). (A,B) As shown by representative traces, exposure to calcium triggered hTRPA1 and Δ1-688 hTRPA1 outward single-channel currents of various magnitude and frequency. (C) Traces showing inhibition of calcium-evoked hTRPA1 activity by the TRPA1 antagonists A-967079 and HC030031 (n=3). (D) Calculated single-channel open probability and conductance values. Data is from 4-6 separate experiments. C = closed-channel state, upward deflection = open channel state.

## Discussion

The effect of calcium on TRPA1 is mechanistically intriguing, and encompasses channel activation by calcium alone as well as calcium channel sensitization/desensitization of ligand activation (5). These effects of calcium may occur by a direct interaction with TRPA1 or as a result of association with a calcium binding partner such as calmodulin (6-9, 11-14). The site of action may be on the cytoplasmic N- and C-termini (5). In this, study, we have addressed the possibility of a direct interaction between calcium and purified hTRPA1 without any interplay with other calcium-sensitive proteins including calmodulin, TRPV1 and A-kinase anchoring protein (AKAP) that can associate with TRPA1 and influence its function (5, 13).

As shown in FRET experiments, the FRET efficiency distributions of both hTRPA1 and Δ1-688 drastically changed in the presence of 200 µM calcium, indicating structural rearrangements caused by a direct interaction between calcium and channel structures outside the N-ARD. The FCS experiments confirmed that all samples contained the same number of fluorescently-labeled hTRPA1 molecules in the confocal volume under the various experimental conditions. Furthermore, when 2 µM of non-labeled hTRPA1 proteins were added to 50 nM of labeled hTRPA1 proteins, the same fluorescence brightness and similar hydrodynamic radii were observed as in the absence of non-labeled TRPA1 proteins. These results indicate that *in vitro* there is no substantial shuffling between the labeled and non-labeled hTRPA1 monomers, and thus both hTRPA1 and Δ1-688 hTRPA1 exist as a stable tetramer complex in the aqueous test solution. Although it could be argued that the changes in FRET efficiency induced by calcium is due to the formation of TRPA1 aggregates formed by intermolecular interactions (21), the fluorescence intensity traces showed no signs of aggregates. Interestingly, in the presence of calcium, both hTRPA1 and Δ1-688 hTRPA1 showed a similar donor-acceptors global distance of 79 Å and 76 Å, respectively, suggesting that intramolecular repulsions were the same in both TRPA1 proteins. This may suggest that calcium binds to the same sites in hTRPA1 and Δ1-688 hTRPA1.

To get further insight into the functional consequences of calcium-induced structural changes of hTRPA1 with and without its N-ARD, we reconstituted the purified hTRPA1 proteins into artificial lipid bilayers for studies of electrical activity as in previous studies of hTRPA1 intrinsic chemo-thermo- and mechanosensitivity (17-20). At calcium concentrations that evoked half and near maximal activation of heterologously expressed TRPA1 in isolated inside-out cell membrane patches (6-8), we recorded intense activity of both hTRPA1 and Δ1-688 hTRPA1. There was no obvious difference with regard to single-channel open probability and conductance levels between the hTRPA1 proteins. However, compared to activation of purified hTRPA1 by most ligands and temperature (17-19), using the same experimental set-up and conditions, calcium seemed to evoke more variability in open probability and conductance levels of both hTRPA1 proteins. Thus, in a physiological context, there must be a need for intracellular co-factors such as calmodulin to tightly control and fine-tune the otherwise wild activity of TRPA1 triggered by calcium (9, 13). Importantly, in a mass spectrometry study on hTRPA1 and Δ1-688 hTRPA1 (22), we did not find any calcium binding partners including calmodulin associated with hTRPA1 and Δ1-688 hTRPA1, which could have been reminiscent of the expression and purification of the hTRPA1 proteins although performed in *Pichia pastoris*. Our aim was not to study calcium sensitization/desensitization properties of TRPA1 by calcium alone or in the presence of TRPA1 activators such as cinnamaldehyde and AITC, but we noticed a lack of calcium self-desensitization in recordings even up to a minute, which is in contrast to recordings in excised inside-out patches within the same timeframe (7). As the hTRPA1without its intracellular C-terminus could not be expressed, the possibility that calcium also modulates hTRPA1 activity by interacting directly with a putative EF-hand calcium-binding domain in the N-terminus cannot be excluded. Nevertheless, our findings that calcium interacts with TRPA1 outside the N-ARD is in line with studies suggesting specific calcium binding sites in the C-terminus and/or in the cytoplasmic end of transmembrane domain 2 and 3 of TRPA1 (11, 13, 14).

In conclusion, calcium directly interacts with hTRPA1 outside its N-ARD in a non-calmodulin-dependent manner, causing structural changes and TRPA1 channel activity.

## Materials and Methods

### Recombinant protein expression and purification

The hTRPA1 and Δ1-688 hTRPA1 were expressed in *Pichia pastoris* and purified as described earlier (19). The purity of recombinant proteins was checked by SDS/PAGE followed by Coomassie Blue R-250 staining and mass spectrometry. The concentrations of all proteins were calculated using Bradford assays. For all experiments, His-tagged recombinant hTRPA1 proteins were used as it was shown that wild-type His-tagged hTRPA1 behaves like the native protein, indicating that the His-tag does not interfere in our experiments (19).

### Protein labeling

In FCS experiments, the hTRPA1 and the Δ1-688 hTRPA1 N-terminal-His-tag was labeled with the Ni-NTA (Nα,Nα-bis(carboxymethyl)-L-lysine, Nickel(II) Atto647N (Sigma-Aldrich) following the protocol provided by the manufacturer. In FRET experiments, besides the lysine labeling with acceptor Atto647N (Atto-Tech), the N-terminal-His-tag was labeled with the Ni-NTA Atto550 fluorophore (Sigma-Aldrich) following the protocol provided by the manufacturer. In all cases, excess fluorophore was added to the aqueous protein solution containing 50 mM PBS buffer at pH 7.8 supplemented with 130 mM NaCl and 0.014% phosphatidyl choline (FC14) (Sigma) and left for 8 h at 4°C. The labeled protein was subsequently purified using size exclusion chromatography on a Sephadex™ G-25 Medium column (GE Healthcare, UK). To remove excess free dyes from the labeled proteins and also avoid hydrolysis of extra fluorophore, the samples were dialysed (3k membrane) for two times, four hours of incubation in each time at 4°C. For long storage, the conjugates were divided into small aliquots and frozen at 20°C to avoid repeated freezing and thawing.

### Sample preparation

In FCS experiments, the concentration of the Atto647N labeled hTRPA1 and Δ1-688 hTRPA1 in the measuring buffer was 50 nM and 2 mM of CaCl_2_, both of which were incubated together for 2 h at 22°C. In FRET experiments, the concentration of the Atto550/Atto647N double-labeled hTRPA1 and Δ1-688 hTRPA1 was 10 nM and 200 µM of CaCl_2_, both of which were incubated together for 2 h at 22°C. The freshly prepared samples were immediately used for FCS and FRET experiments. In both assays, 0.1% Tween-20 (Sigma) was added in order to diminish surface interactions with the glass coverslip.

### Fluorescence correlation spectroscopy (FCS)

The FCS experiments were carried out at 22°C with a custom-built confocal fluorescence microscope with a Zeiss C-Apochromat 40 × 1.2 NA water-immersion objective. The fluorescence intensity temporal fluctuations were analyzed with a hardware correlator (Flex02-12D/C correlator with 12.5 ns minimum channel width). All the experimental data were fitted by considering a single species and free Brownian 3D diffusion in the case of a Gaussian molecular detection efficiency:

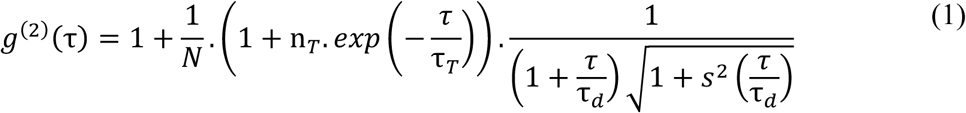

where N is the average number of molecules in the focal volume, n_T_ is the amplitude of the dark state population, τ_T_ is the dark state blinking time, *τ*_*d*_ is the mean diffusion time and s is the ratio of transversal to axial dimensions of the analysis volume. The molecular diffusion coefficient, D and hydrodynamic radius, R_H_ were calculated as previously as described in detail (23). The confocal volume was defined by using the 30 μm confocal pinhole conjugated to the sample plane whose transversal waist *w*_*xy*_ was calibrated to 285 nm using the known diffusion coefficient of Alexa 647 in pure water (3.1 × 10^−6^ cm^2^·s^−1^ at 22°C) and known hydrodynamic radius of 0.7 nm in pure water. Each FCS measurement lasted for 100 seconds, and measurements were repeated several times on different days to determine the average translational diffusion times.

### Förster resonance energy transfer (FRET)

The FRET signal was detected using a confocal inverted microscope with a Zeiss C-Apochromat 63×1.2 NA water-immersion objective, and an iChrome-TVIS laser (Toptica GmbH) as an exciting source operating at 550 nm. Filtering the laser excitation was achieved by a set of two bandpass filters (Chroma ET525/70M and Semrock FF01-550/88). Dichroic mirrors (Chroma ZT594RDC and ZT633RDC) separate the donor and acceptor fluorescence light. The excitation power at the diffraction limited spot was set to 20 μW for all experimental conditions. The detection was performed by two avalanche photodiodes (Micro Photon Devices MPD-5CTC with 50 μm active surface) with 620 ± 20 nm (Chroma ET605/70M and ET632/60M) and 670 ± 20 nm (Semrock FF01-676/37) fluorescence bandpass filters for the donor and acceptor channels respectively. The photodiode signal was recorded by a fast time-correlated single photon counting module (Hydraharp400, Picoquant GmbH) in time-tagged time-resolved (TTTR) mode. Conceptually, the apparent FRET efficiency of each burst was calculated as previously described in detail (24). All fluorescence bursts above the background noise were recorded separately by the acceptor channel and donor channel. As a part of calibration, we took into consideration the differences in the fluorescence detection efficiencies (η_A_ and η_D_), direct excitation of the acceptor by the laser light 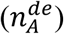, donor emission crosstalk into the acceptor channel (α) and quantum yields of fluorophores (φ_A_ and φ_D_). Symphotime 64 (Picoquant GmbH) software was used to compute the FRET efficiency according to the formula below:

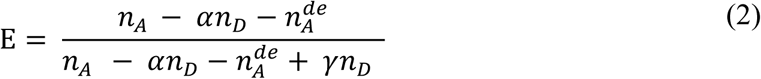

where γ = η_A_φ_A_/η_D_ φ_D_ accounts for the differences in quantum yields (φ_A_ and φ_D_) and fluorescence detection efficiencies (η_A_ and η_D_) between the acceptor and donor.

We estimate γ = 1.3, and α = 0.16 for the current setup.

### Planar lipid bilayer patch-clamp electrophysiology

These experiments were performed as previously described in detail (19) and are briefly described as follows. Purified hTRPA1 was reconstituted into preformed planar lipid bilayers composed of 1,2-diphytanoyl-sn-glycero-3-phosphocholine (Avanti Polar Lipids) and cholesterol (Sigma-Aldrich) in a 9:1 ratio and produced by using the Vesicle Prep Pro Station (Nanion Technologies). Under these conditions, a uniform protein orientation is favored with N- and C-termini facing the recording chamber (i.e., the “cytosolic compartment”). Ion channel activity was recorded using the Port-a-Patch (Nanion Technologies) at a positive test potential of +60 mV in a symmetrical K^+^ solution (50 mM KCl, 10 mM NaCl, 60 mM KF, 20 mM EGTA, and 10 mM Hepes; adjusted to pH 7.2 with KOH) and at room temperature (20-22°C). To prepare free Ca^2+^ solutions, the concentrations of calcium were calculated with MaxChelator (https://somapp.ucdmc.ucdavis.edu/pharmacology/bers/maxchelator/downloads.htm). Signals were acquired with an EPC 10 amplifier and PatchMaster software (HEKA) at a sampling rate of 50 kHz. Electrophysiological data were analyzed using Clampfit 9 (Molecular Devices) and Igor Pro (WaveMetrics). Data were processed by a Gaussian low-pass filter at 1000 for analysis and 500 Hz for traces. The single-channel open probability (P_o_) was calculated from time constant values, which were obtained from exponential standard fits of dwell time histograms. The single-channel conductance (G_s_) was obtained from Gaussian fit of all-points amplitude histograms.

## Acknowledgements

This study was supported by the Swedish Research Council (2014-3801) and the Medical Faculty of Lund University - ALF (Dnr. ALFSKANE-451751). This project has received funding from the Agence Nationale de la Recherche (ANR) under grant agreement ANR-17-CE09-0026-01 and from the European Research Council (ERC) under the European Commission’s Seventh Framework Programme (grant agreement 278242).

The authors declare no competing interest.

